# Independent benchmark of H&E-based gene expression prediction in skin

**DOI:** 10.64898/2026.09.07.749926

**Authors:** Regina Shaikhutdinova, Sabina Gansberger, Julia Staller, Namrata Singh, Inigo Oyarzun, Martin Simon, Barbara Sterniczky, Philipp Tschandl, Johannes Griss

**Author notes:** **Correspondence:** Johannes Griss, Medical University of Vienna Department of Dermatology Währinger Gürtel 18-20, 1090 Vienna, Austria.

## Abstract

Given the widespread availability of H&E slides, there is considerable interest in determining whether molecular information can be inferred directly from tissue morphology, potentially reducing the need for costly spatial transcriptomic profiling. We assessed three state-of-the-art methods for predicting single-cell gene expression from H&E images across three skin disease contexts and two Xenium panels. As controls, we included simple linear regression models trained on embeddings from multiple foundation models, totalling 16 models evaluated in this study. We show that all models performed poorly: for most genes, prediction accuracy was near zero, and reliable predictions were largely restricted to keratinocyte-associated genes. Predicted expression failed to preserve cell-type identity and spatial organisation, with only keratinocytes forming coherent clusters, while immune, fibroblast, and other dermal populations were extensively mixed. Notably, simple ridge regression on pretrained embeddings matched or outperformed the more complex published architectures, indicating that the predictive signal originates primarily from image representations rather than model design. Our results demonstrate that current H&E-based gene expression prediction methods are not yet suitable for single-cell-level interpretation of spatial transcriptomics in skin tissue.

**Key points:**

- We provide an independent benchmark of state-of-the-art H&E-based single-cell gene expression prediction methods in human skin.
- Current methods accurately predict only a limited subset of genes, predominantly those associated with keratinocytes.
- Complex deep learning architectures provide little improvement over simple linear models built on pretrained image embeddings.
- Biological cell-type structure and spatial organization are not reliably preserved in predicted expression profiles.

## Introduction

H&E-stained tissue sections are widely used in clinical pathology due to their ability to provide rapid, reliable, and cost-effective assessment of tissue architecture and cellular organization (Fischer *et al.* 2008). Recent advances in machine learning have substantially expanded the utility of H&E images, enabling representation learning, cross-modal alignment, and the prediction of spatial omics directly from histological data (Hao *et al.* 2026).

The emergence of high-resolution spatial transcriptomics platforms, such as 10X Xenium, has further accelerated this field by providing single-cell, spatially resolved gene expression measurements that can serve as ground truth for training and evaluating predictive models. In parallel, large pathology foundation models trained on extensive histology datasets have produced powerful image representations (He *et al.* 2015, Filiot *et al.* 2023, Chen *et al.* 2024, Xu *et al.* 2024, Hemker *et al.* 2026), raising the prospect that gene expression might be inferred directly from routine H&E sections. If accurate, such predictions could reduce the cost and complexity of spatial profiling.

Despite these advances, comprehensive independent benchmarking studies remain limited (Zhu *et al.* 2026). Here, we present a benchmark of state-of-the-art models for predicting gene expression from H&E images at single-cell resolution in human skin samples. By evaluating multiple datasets and disease contexts, we assess the extent to which current approaches can accurately reconstruct spatial transcriptomic information in skin tissue.

## Results

We evaluated three recently published methods (Figure 1a): GHIST (Fu *et al.* 2025), a deep learning framework based on a UNet3+ backbone trained from scratch on paired H&E and spatial transcriptomics data; Pixel2Gene, which uses the HIPT pathology foundation (Zhang *et al.* 2024) model to extract histological features that are subsequently processed by a simple multilayer perceptron (MLP); and SpatialEx (Liu *et al.* 2026), a hypergraph-based model trained on embeddings generated by the UNI (Chen *et al.* 2024), Phikon (Filiot *et al.* 2023), GigaPath (Xu *et al.* 2024), ResNet50, ResNet101, and ResNet152 (He *et al.* 2015) pretrained image encoders, all of which are natively supported by SpatialEx. In addition, we integrated the SEAL foundation model (Hemker *et al.* 2026), which unlike the other foundation models, was trained on both H&E images and spatial transcriptomics data into SpatialEx. As controls, we trained simple linear ridge regression models that take the same pretrained image-encoder embeddings as input and directly predict gene expression. Ridge regression models were trained on embeddings from each of the same encoders used by SpatialEx, resulting in a total of 16 models evaluated in this study (Figure 1a).

**Figure 1.**
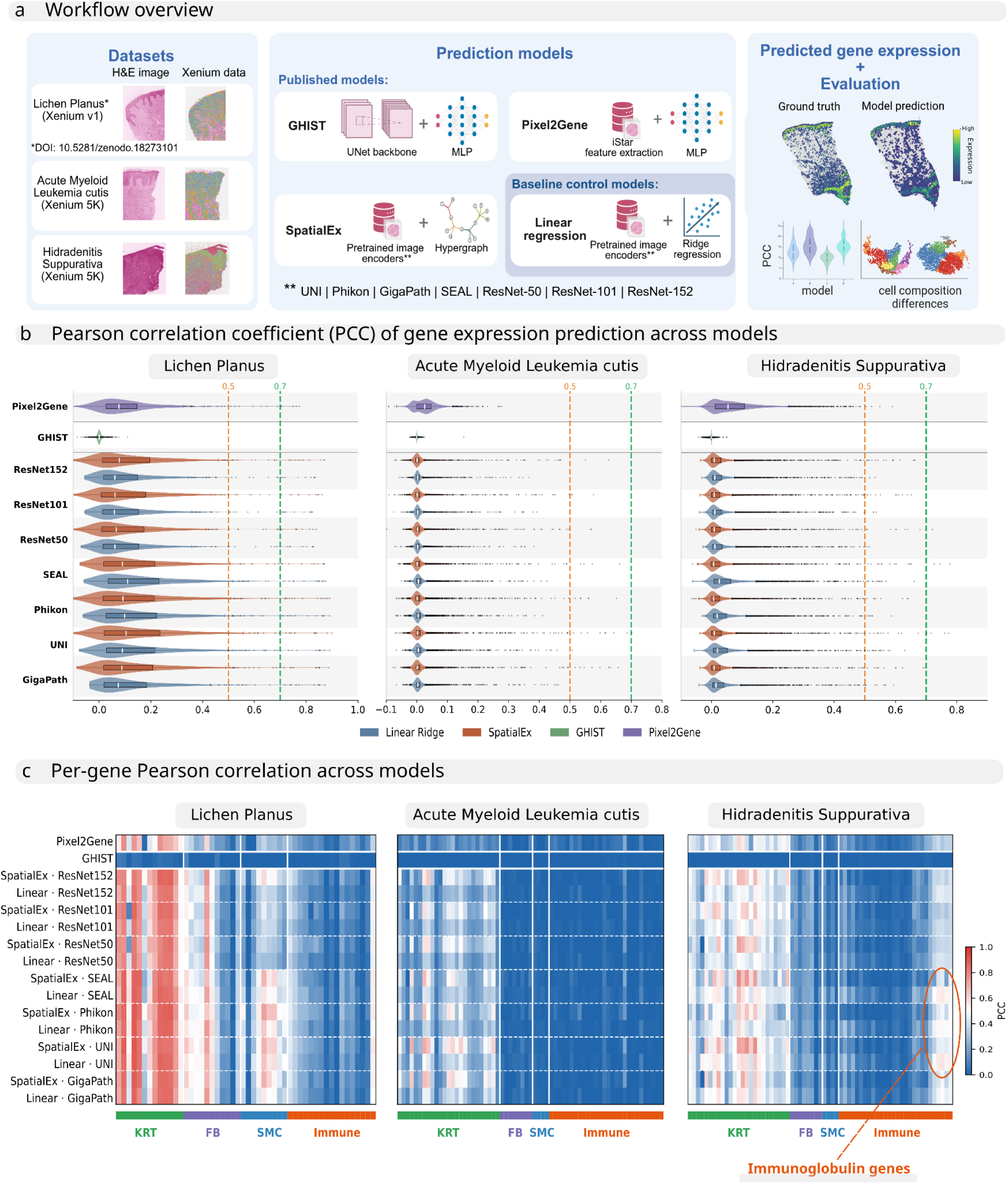
Performance of benchmarked models on three skin tissue datasets. **(a)** Graphical summary of the study’s workflow. **(b)** Per-gene Pearson correlation coefficient (PCC) per model and dataset. Dashed lines mark PCC 0.5, 0.7. **(c)** Per-gene PCC for grouped cell-type markers: keratinocyte- (KRT), fibroblast- (FB), smooth muscle cell- (SMC), and immune-associated genes (Immune). **Alt text:** Four-panel figure benchmarking gene-expression-from-histology models. Panel a is a workflow schematic: H&E images and matched Xenium spatial transcriptomics from three skin datasets (hidradenitis suppurativa, AML cutis, lichen planus) feed three published methods (GHIST, Pixel2Gene, SpatialEx) and linear ridge regression baselines; predictions are compared to ground truth using PCC, SCC and SSIM. Panel b is a grid of per-gene PCC distributions (violin plots) by model and dataset, with dashed lines at 0.5 and 0.7. Panel c is a heatmap of per-gene PCC for markers grouped as keratinocyte, fibroblast, smooth muscle and immune.

Each model was trained and validated on a tissue section where both Xenium data and the H&E image were available. Training and validation slides were from separate individuals and predicted gene expression was compared to the Xenium ground truth. All models were evaluated using in-house Xenium 5K panel data generated from hidradenitis suppurativa (HS) and acute myeloid leukemia cutis (AML cutis) skin tissue samples, together with a publicly available Xenium v1 dataset of Lichen Planus (LP) (Dey 2026). Model performance was assessed using three different metrics (Figure 1b, Supplementary Figure 1a, b): the Pearson correlation coefficient (PCC), the Spearman correlation coefficient (SCC), and the Structural Similarity Index Measure (SSIM).

### Current methods fail to reliably predict the majority of genes in skin datasets

SpatialEx generally outperformed GHIST and Pixel2Gene in terms of the mean PCC and number of genes achieving a PCC greater than 0.5 (Figure 1b, Supplementary Figure 1c). However, the majority of genes showed a mean PCC value close to zero, and only 5.4% of genes achieved a PCC above 0.5 in the Xenium v1 LP dataset for the best-performing model. In the Xenium 5K HS dataset, the corresponding value was only 0.3% (Supplementary Figure 1c). Interestingly, the LP dataset contained a greater number and proportion of genes reaching PCC > 0.5, as well as higher mean PCC values overall. Reducing the HS dataset from the 5K gene panel to the v1 panel did not improve performance at the individual-gene level (Supplementary Note 1). This difference may therefore reflect the higher per-transcript sensitivity of the v1 panel, or differences in tissue composition and disease context between the datasets, rather than the smaller panel size alone.

Surprisingly, simple ridge regression models trained on the same image embeddings achieved a performance comparable to or even exceeding SpatialEx (Figure 1b, Supplementary Figure 1c). In the HS dataset, the best-performing linear ridge model (selected based on mean PCC and the number of genes with PCC > 0.5) achieved a mean PCC of 0.043, compared to 0.03 for the best-performing SpatialEx model. In the LP dataset, SpatialEx slightly outperformed linear ridge regression (mean PCC 0.158 vs. 0.151). This finding suggests that the predictive information is already captured by the pretrained image encoders, with only limited additional benefit of a more complex architecture.

### Reliable gene prediction is largely restricted to keratinocyte-associated genes

We next assessed whether prediction accuracy differed between genes (Figure 1c). For interpretation, genes were grouped into four major categories: keratinocyte-associated (KRT), fibroblast-associated (FB), smooth muscle cell-associated (SMC), and immune-related (Immune) genes.

For the HS dataset, prediction performance was consistently highest for keratinocyte-associated genes, likely reflecting the strong correlation of epithelial morphology with keratinocyte transcriptional programs (Figure 1c). In contrast, genes associated with fibroblasts, smooth muscle cells, and immune cell populations were generally predicted poorly. Notably, immunoglobulin genes, including IGHG1, IGHG2, IGHG3, IGHG4, and IGKC, represented an exception and achieved comparatively high prediction accuracy in the HS dataset (average PCC 0.54, Figure 1c, Supplementary Table 1). This may reflect the distinctive histomorphological features and increased abundance of plasmablasts/plasma cells in HS tissue. In contrast, these genes were not predicted successfully in the AML dataset (average PCC 0.005, Supplementary Table 2).

Overall, prediction accuracy was higher in the LP dataset, particularly for keratinocyte-associated genes, as well as subsets of smooth muscle cell and fibroblast genes. However, prediction performance for immune-associated genes remained poor.

Spatial maps of gene expression for genes from different cell-type categories further confirmed these results. KRT5 was predicted with sufficient accuracy (Figure 2a; PCC for the best-performing model = ∼0.77), reflected by matching expression patterns in the visualization. Nevertheless, PDGFRA expression prediction largely failed even with the best-performing model: the highest accuracy was achieved by linear ridge UNI (PCC =∼ 0.13), while the best SpatialEx model, SpatialEx ResNet101, reached only PCC =∼ 0.08, despite this gene being among the top 15% most highly expressed transcripts in the ground truth dataset and predominantly expressed by fibroblasts in skin tissue. This low accuracy is reflected by predicted expression patterns that do not match the ground truth (Figure 2b). A similar pattern was observed for CD19 (Figure 2c; PCC =∼ 0.17 for linear ridge SEAL, and PCC =∼ 0.15 for SpatialEx ResNet152), which is among the top 25% most highly expressed transcripts in the ground truth dataset and is specific to B cells. Overall, this suggests that successful prediction depends on the presence of a spatial pattern that corresponds to gene expression, such as the epithelial localization of keratins.

**Figure 2.**
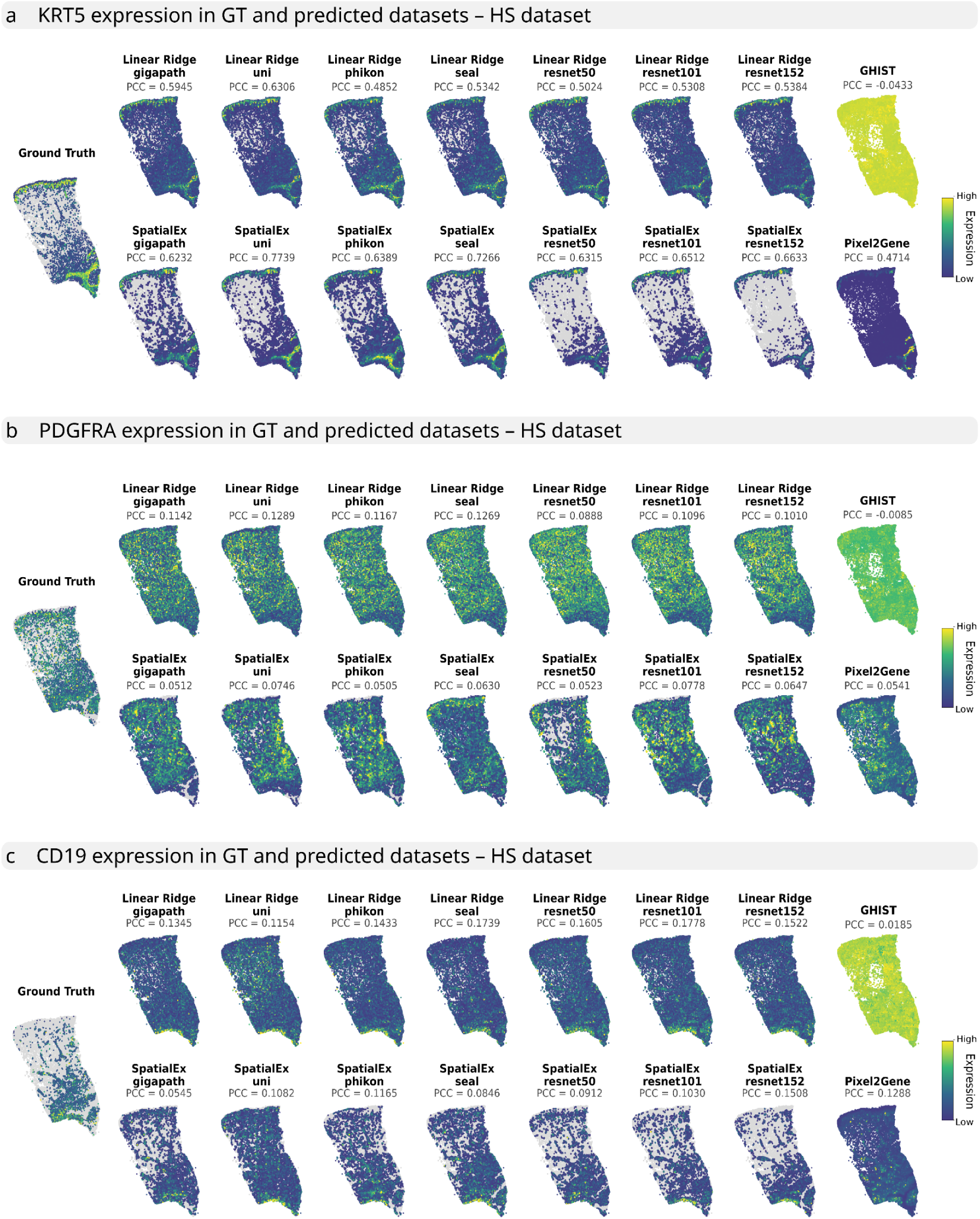
Spatial visualization of predicted and measured expression for three representative genes. **(a)** Expression of KRT5; the best-performing model was SpatialEx UNI (PCC = 0.7739). **(b)** Expression of PDGFRA; the best-performing model was Linear Ridge UNI (PCC = 0.1289). **(c)** Expression of CD19; the best-performing model was Linear Ridge ResNet101 (PCC = 0.1778). In all panels, color represents expression level, with yellow indicating high and dark blue indicating low expression. **Alt text:** Grid of spatial plots comparing predicted versus measured single-cell expression for three genes across models, with cells colored by expression. Panel a (KRT5, keratinocyte) shows predicted maps that closely match the measured epithelial pattern. Panel b (PDGFRA, fibroblast) and panel c (CD19, B cell) show predicted maps that fail to reproduce the measured spatial pattern, appearing diffuse or near-uniform.

### Limited preservation of biological cell-type structure in predicted gene expression in HS dataset

To assess whether predicted gene expression preserves biological cell-type structure, we focused on the HS dataset and selected SpatialEx UNI and linear ridge UNI as the two best-performing models (Supplementary Figure 1c). We visualized their predictions using UMAP embeddings and spatial coordinates (Figure 3a).

**Figure 3.**
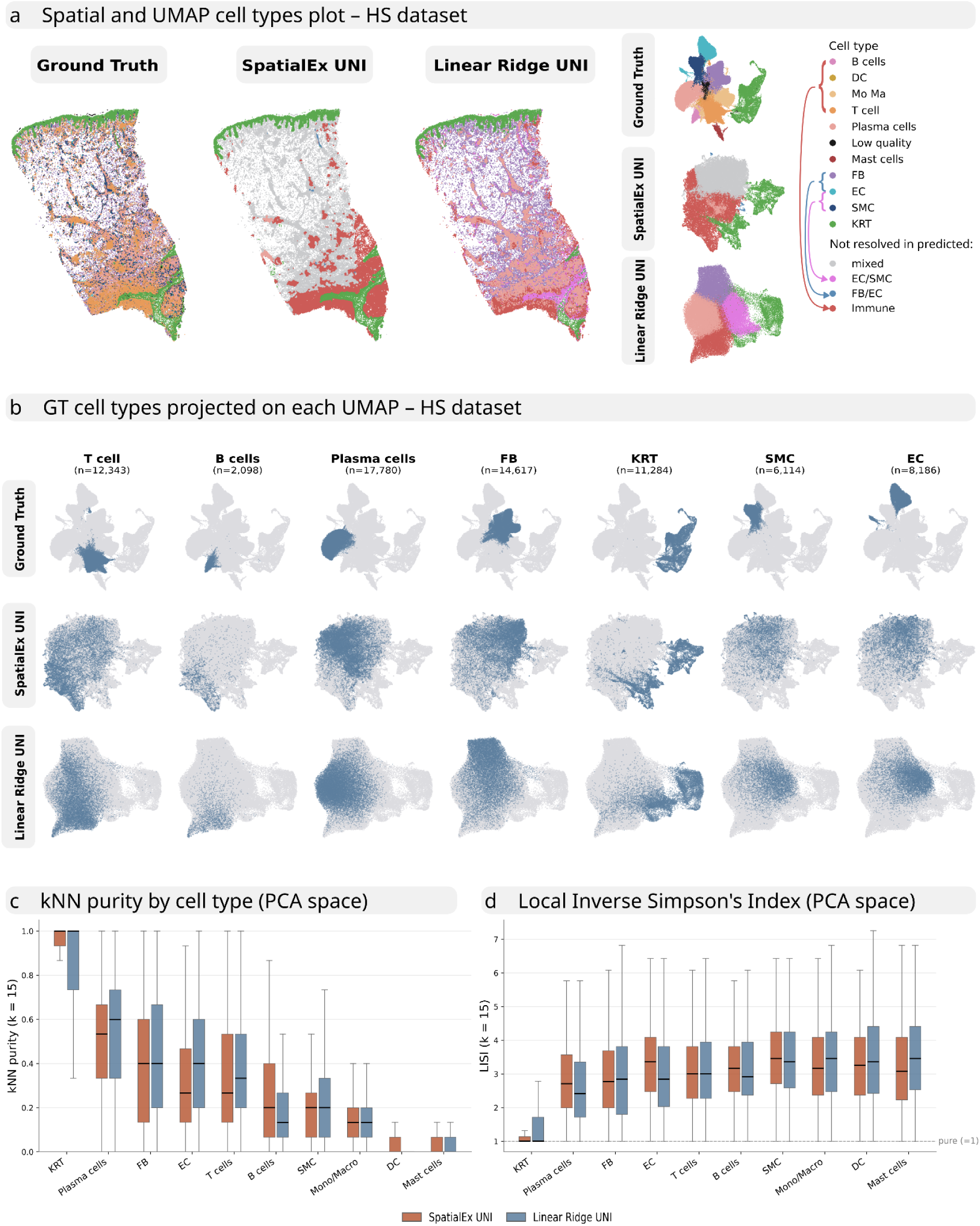
Cell-type structure in predicted gene expression, HS dataset. **(a)** Spatial and UMAP visualization of ground truth and predicted cell types in HS for the two best models. Colors represent annotated cell types. **(b)** Ground truth labels projected onto each UMAP. **(c)** kNN purity by cell type (PCA space); values near 1 indicate higher purity. **(d)** Local Inverse Simpson’s Index (LISI) by cell type (PCA space); LISI = 1 indicates purity, higher values mixing. **Alt text:** Figure on cell-type structure in the HS dataset for ground truth versus the two best models (SpatialEx UNI, Linear Ridge UNI). Panel a shows paired spatial plots and UMAP embeddings colored by annotated cell type. Panel b overlays ground-truth labels onto each predicted UMAP. Panel c is a bar/box plot of kNN purity by cell type. Panel d is a plot of LISI by cell type.

Clusters were annotated using canonical marker genes (Supplementary Figure 2a, b). For the predicted datasets, most clusters could not be confidently annotated, as they exhibited mixed expression signatures of multiple cell types, therefore all annotations for the predicted datasets should be considered approximations. In particular, fibroblasts and endothelial cells merged into a single cluster in the SpatialEx predictions, while endothelial cells and smooth muscle cells merged into a single cluster in the linear ridge predictions (Figure 3a). In both methods, B cells, T cells, macrophages, monocytes, and dendritic cells collapsed into a single immune cluster (Figure 3a, b). Additionally, the SpatialEx predictions contained a mixed cell population expressing markers of fibroblasts, endothelial cells, and immune cells simultaneously (Figure 3a).

Interestingly, based on the cluster-level gene expression profiles (Supplementary Figure 2a, b), ridge regression was able to partially resolve fibroblast and plasma cell clusters, whereas SpatialEx failed to separate these populations entirely (Figure 3a). However, both methods produced substantially more disorganized embeddings than the ground truth dataset. Neither model was able to distinguish immune subtypes, with the exception of plasma cells, which were partially resolved in the ridge regression predictions. In the Linear Ridge predictions, plasma cells were better resolved and the spatial organization of the dermal layer more closely resembled the ground truth than in the SpatialEx predictions (Figure 3a).

To measure how well the predicted data preserves biological structure, we projected ground truth cell-type labels onto the SpatialEx and Linear Ridge embeddings by shared cell index (Figure 3b). For all cell types except keratinocytes, the ground truth labels were dispersed across the predicted UMAPs rather than forming distinct groups, indicating that the predictions do not preserve the biological structure of the dataset.

To quantify this, we used two complementary metrics. First, we calculated kNN purity (k = 15) in PCA space (Figure 3c). Although the predicted embeddings recovered keratinocytes as a relatively pure cluster (kNN purity above 0.8), purity was substantially lower for the other cell types (below 0.5), confirming that cell-type structure is poorly preserved overall. Second, we calculated the local inverse Simpson’s index (LISI; Figure 3d). LISI confirmed that only keratinocytes maintained pure neighborhoods in the predicted PCA space, whereas all other cell types showed substantial mixing, with neighborhoods containing at least two cell types on average (median LISI > 2).

These findings indicate that current models are not yet suitable for single-cell level interpretation of spatial transcriptomics predictions in skin, as substantial noise remains and many cell types cannot be reliably distinguished.

### Limited preservation of biological cell-type structure in predicted gene expression in LP dataset

We repeated the same cell-level analysis on the Lichen Planus v1 Xenium dataset. As before, we used the best-performing models, SpatialEx UNI and linear ridge UNI (Supplementary Figure 1c). Again, the models failed to distinguish between different immune cell types (Figure 4a). SpatialEx produced a large cluster of mixed cells that could not be annotated due to the shared expression of markers from multiple cell types (Supplementary Figure 4), recovering only keratinocytes and a mixed immune cluster. Linear ridge regression performed slightly better, recovering keratinocytes, fibroblasts, smooth muscle cells, and a mixed cell cluster. We observed that ground truth labels when projected onto the predicted UMAPs (Figure 4b) appeared highly dispersed. Consistent with this, kNN purity was high only for keratinocytes (above 0.8) and fell below 0.5 for all other cell types (Figure 4c), and LISI was 1 only for keratinocytes (Figure 4d). These results show that the poor prediction of non-keratinocyte cell types persists even on v1 Xenium panel.

**Figure 4.**
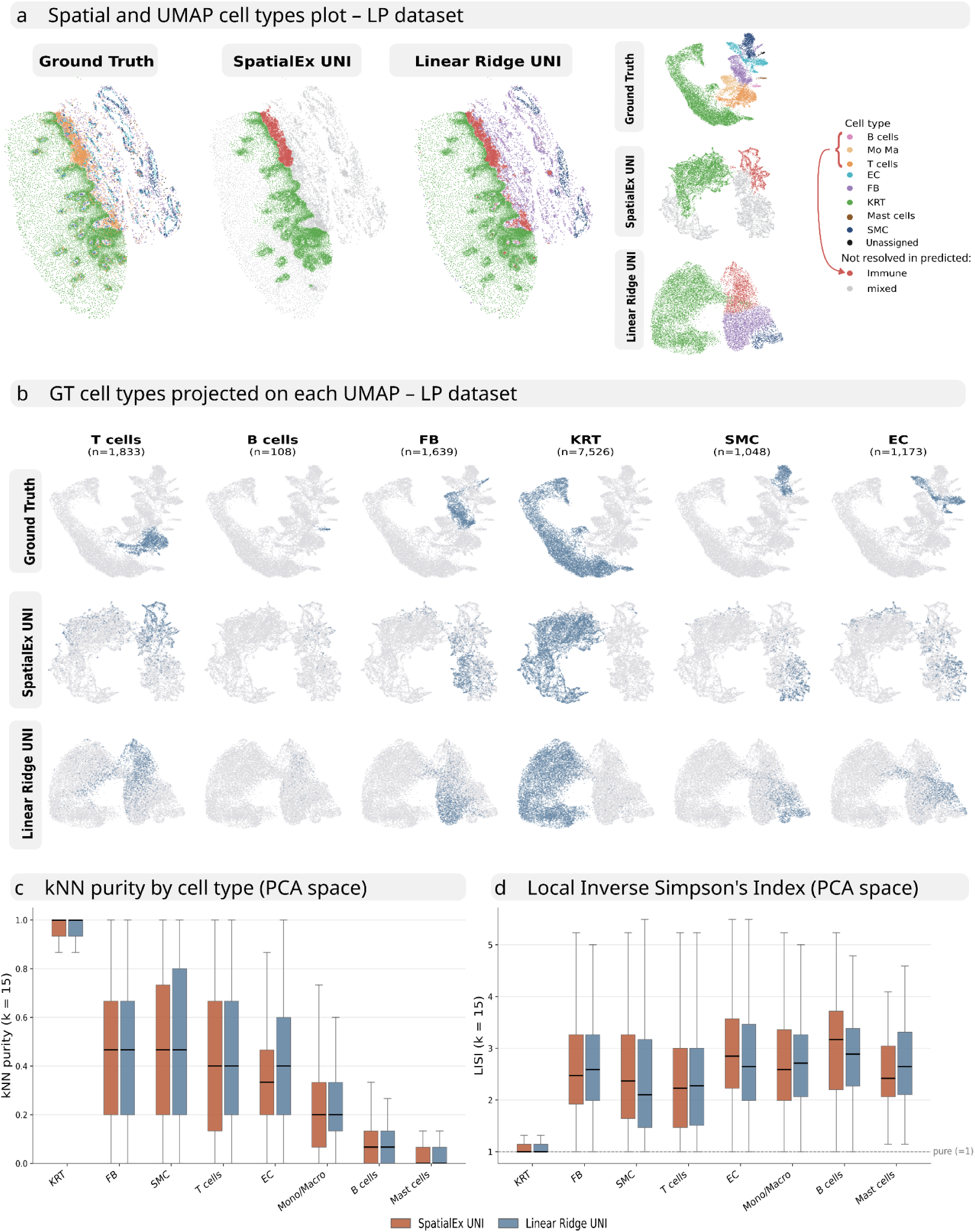
Cell-type structure in predicted gene expression, LP dataset. **(a)** Spatial and UMAP visualization of ground truth and predicted cell types in LP for the two best models. Colors represent annotated cell types. **(b)** Ground truth labels projected onto each UMAP. **(c)** kNN purity by cell type (PCA space); values near 1 indicate higher purity. **(d)** Local Inverse Simpson’s Index (LISI) by cell type (PCA space); LISI = 1 indicates purity, higher values mixing. **Alt text:** Same as Figure 3 but for the LP dataset. Panel a: spatial and UMAP plots colored by cell type. Panel b: ground-truth labels projected onto predicted UMAPs. Panel c: kNN purity. Panel d is a plot of LISI by cell type.

## Discussion

Given the rapid pace of development in the field, benchmarking the most recent methods is challenging, which is why previous benchmarks have largely focused on spot-based and generally older methods (Xie *et al.* 2023, Wang *et al.* 2025). To our knowledge, no prior study has compared these recent, more complex models against simple linear regression trained on embeddings from different pretrained encoders. Additionally, several reviews have highlighted the need for standardized, independent benchmarking of these models (Hao *et al.* 2026, Zhu *et al.* 2026). Such independent evaluation is especially important because predictions that appear accurate can still contribute little biological insight (Hao *et al.* 2026).

The poor performance of all models contrasts the original publications, and highlights potential issues in benchmarking predictive methods for spatial transcriptomics. For example, SpatialEx was developed and evaluated on consecutive sections from the same 10x Genomics Xenium FFPE breast cancer specimen (Janesick *et al.* 2023, 10x Genomics 2024). Pixel2Gene’s prediction of gene expression from H&E alone was supported solely by qualitative visualizations of selected marker genes, without quantitative comparison to a measured ground truth (Yao *et al.* 2026). In our benchmark, we instead evaluated all models on tissue sections from different individuals, a more demanding cross-patient design that requires generalization to unseen morphological and molecular variation rather than reproduction of a near-identical replicate.

Additionally, for SpatialEx, the authors focused primarily on well-predicted genes (Liu *et al.* 2026). Similarly, for GHIST the authors emphasized performance on the top 20 highly variable genes (Fu *et al.* 2025).

Our results reveal that all evaluated methods performed poorly on skin tissue when applied to truly unseen samples: for most genes, predicted and measured expressions were essentially uncorrelated, and only a small fraction reached a PCC above 0.5. Reliable prediction was largely restricted to keratinocyte-associated genes, likely reflecting the close coupling between epithelial morphology and keratinocyte transcriptional programs, as keratinocytes occupy a spatially distinct, morphologically unambiguous compartment that is readily recognizable in H&E (Yousef *et al.* 2026). In contrast, genes associated with fibroblasts, smooth muscle cells, and immune populations were predicted poorly, as these cell types are interspersed within the dermis and lack distinctive morphological signatures. The main exception were immunoglobulin genes, predicted comparatively well in the HS dataset, reflecting the distinctive morphology and high local abundance of plasma cells in hidradenitis suppurativa. Moreover, predicted expression did not preserve biological phenotypes, with only keratinocytes forming pure clusters while immune, fibroblast, and other dermal populations collapsed together.

Taken together, current H&E-based prediction methods are not yet suitable for single-cell-level interpretation of spatial transcriptomics in skin. Reliable prediction is confined to morphologically distinctive keratinocyte-associated genes, while the dermal and immune compartments central to most inflammatory and neoplastic skin diseases are not faithfully recovered. Until predicted expression can preserve cell-type identity and spatial organization across these compartments, it should not substitute for direct measurement.

## Methods

### Xenium in-house data

Formalin-fixed paraffin-embedded (FFPE) samples from clinical routine samples fixated after biopsy with 7.5% formaldehyde and tissue sections were profiled using the Xenium Prime 5K Human Pan-Tissue and Pathways panel (10x Genomics), supplemented with a custom panel of 51 additional genes.

Tissue sections were cut at a thickness of 5 μm following the Xenium In Situ for FFPE-Tissue Preparation Guide (CG000578, 10x Genomics), spread out in RNA enzyme-free water at 38 °C, and attached to the Xenium slides (PN-3000941, 10x Genomics) within the sample area. The slides were dried at room temperature for 30 min and baked at 42 °C for 3 h. The follow-up experiment was carried out after drying for five days at room temperature. All further steps were carried out according to the Xenium Prime In Situ Expression protocol (CG000760, 10x Genomics).

The slides were processed on the Xenium Analyzer instrument for in situ transcriptomic profiling. Subsequently, cell segmentation was carried out using nuclear staining and membrane markers to define cell boundaries. Transcript assignments, cell segmentation boundaries, and clustering results were visualized using the Xenium Explorer software. Following the Xenium assay, Hematoxylin and Eosin (H&E) staining was performed to assess tissue morphology.

### Public datasets

Xenium spatial transcriptomics data of lichen planus skin lesions were obtained from Zenodo (Dey, P. Lichen_Planus, 2026; DOI: 10.5281/zenodo.18273102; CC-BY 4.0). Tissue sections were profiled using the Xenium Human Skin v1 gene panel (480 genes), with matched post-Xenium H&E histology acquired on the same sections. From sample 10543-JF-1, two regions of interest were used: region D2 was used for model training and region D1 for prediction.

### Published models, training setting and output treatment

#### SpatialEx

During slide preprocessing and model training, we followed <u>Tutorial #1</u> provided by the SpatialEx team, using two Xenium slides. The model was trained with the following parameters: resolution = 64, num_neighbors = 7, epochs = 1000, batch_size = 4096, hidden_dim = 512, num_layers = 2, prune = 10,000, and learning rate = 0.001.

The ResNet models used in conjunction with SpatialEx (ResNet50, ResNet101, and ResNet152) were pretrained on the ImageNet-1k dataset at a resolution of 224×224 pixels.

#### SpatialEx + SEAL

SEAL is a recently published foundation model built on top of CONCHv1 or UNIv2 (in this study, we used the CONCHv1-based version) and is further trained on paired H&E images and spatial transcriptomics data. To incorporate SEAL into SpatialEx, we modified the create_ImageEncoder function in the SpatialEx utility module to include a SEAL-specific loading branch. The model was initialized in frozen inference mode with gradient computation disabled.

All modifications were implemented as runtime monkey-patches applied to the SpatialEx.utils and SpatialEx.preprocess modules, preserving compatibility with the original SpatialEx codebase without requiring changes to the installed package.

#### GHIST

The H&E images were registered to the Xenium outputs using the Xenium Explorer software, and the resulting transformation matrix was exported. This matrix was then applied using the SciPy ndimage module to align the H&E image to the corresponding DAPI image. The registered images were saved in OME-TIFF format and used as input for downstream model training and prediction.

Image preprocessing was performed following the pipeline provided by the original authors, using the same parameter settings. Single-cell reference datasets (GEO accession: GSE287791 for AML and GSE175990, GSE220116 and GSE274880 for HS) were used during training. For the LP dataset, GHIST failed during the H&E nuclei segmentation step; therefore, no single-cell reference dataset was used for this analysis.

The model was trained with the following parameters: embeddings = 512, batch size = 16, learning rate = 0.001, overlap = 64, epochs = 500, with average expression aggregation enabled (avgexp = TRUE), and both cell type (celltype = TRUE) and neighborhood information (neighb = TRUE) incorporated. Multiple parameter configurations were benchmarked, and the final selection was based on performance on a held-out test set, using Pearson correlation coefficient (PCC) as the evaluation metric.

#### Pixel2Gene

Spatial alignment between the H&E image and Xenium coordinate space was performed by manually selecting corresponding keypoints in Napari, using the Xenium morphology image as a reference. These keypoints were used to estimate a transformation matrix, which was applied to transform Xenium transcript coordinates into H&E pixel space.

Preprocessing was conducted using the standard Pixel2Gene pipeline, with minor code modifications required to ensure successful execution.

To obtain cell-level gene expression predictions from the superpixel-resolution outputs, Xenium cell boundaries were mapped to the prediction array coordinate space. Cell boundary vertices in micron coordinates were transformed from stage microns to morphology image pixels, and from morphology pixels to H&E image pixels, followed by scaling to account for H&E image downsampling and superpixel binning. Each cell boundary was represented as a polygon, and only polygons overlapping the prediction array were retained.

For each cell polygon, candidate pixels were first identified using a bounding box filter and then refined using a point-in-polygon test based on pixel center coordinates. Each pixel was assigned to at most one cell. Predicted gene expression values were aggregated per cell by computing the mean across all assigned pixels for each gene. Cells without any assigned pixels, primarily those smaller than the 8 µm superpixel size, were excluded from downstream analyses.

#### Ridge linear regression control

To provide a baseline comparison, we implemented a simple ridge regression model. For each encoder, we used the per-cell embeddings generated during the SpatialEx preprocessing step as input features, ensuring that the ridge models and SpatialEx received identical inputs and differed only in the downstream prediction architecture. A separate ridge regression model was trained for each encoder using the scikit-learn (v1.6.1) Ridge implementation with regularization strength α = 1.0. Each model was trained on one tissue section, using the per-cell embeddings as input and the corresponding measured Xenium expression across all genes, and was then applied to the embeddings of a held-out section from a different individual to predict gene expression.

### Evaluation metrics

Prediction performance was assessed per gene using three metrics: the Pearson correlation coefficient (PCC), the Spearman correlation coefficient (SCC), and the structural similarity index measure (SSIM). For each gene, metrics were computed between predicted and measured expression across all cells of the held-out section.

PCC and SCC were calculated using SciPy, comparing the predicted and measured expression vectors of each gene across all cells; PCC captures linear association and SCC monotonic (rank-based) association. Genes with zero variance in either vector were assigned a correlation of zero, as the coefficient is otherwise undefined.

For SSIM, cells were mapped onto a 50 × 50 spatial grid, and the mean expression per bin was computed for both predicted and measured data. Both grids were normalized to [0, 1], and SSIM was calculated using scikit-image (data range = 1.0), jointly evaluating local luminance, contrast, and structure. Predictions were clipped at zero before evaluation.

For the PCC heatmap (Figure 1c), genes were selected based on their relevance for cell-type annotation, together with additional genes that achieved PCC > 0.5 in at least one model-dataset combination (Supplementary Tables 1-3). Three genes (ARG1, IL1RN, XBP1) were excluded because they were expressed across multiple cell types and lacked the specificity required for cell-type annotation.

### Cell-type structure analysis

For both the HS and LP datasets, the ground truth and predicted expression matrices (SpatialEx UNI and Linear Ridge UNI) were processed independently in scanpy (v1.10.3): feature scaling (max value 10), principal component analysis (30 components), neighborhood graph construction (k = 15), UMAP, and Leiden clustering.

To quantify preservation of cell-type structure, we computed two neighborhood-based metrics in PCA space, after projecting ground truth cell-type labels onto each embedding by shared cell index and excluding cells annotated as “Low quality.” For kNN purity, we identified the 15 nearest neighbors of each cell (Euclidean distance) and calculated the fraction sharing the same ground truth label, averaged per cell type. For the local inverse Simpson’s index (LISI), we computed cell-type proportions among each cell’s 15 nearest neighbors and the inverse Simpson’s index (1 / Σp²), reported as the median per cell type. Both metrics used k = 15 and were computed with scikit-learn (v1.6.1) NearestNeighbors and custom code.

### Compute resources

All models were trained on high-performance computing infrastructure. GHIST and Pixel2Gene, were run on a workstation equipped with an NVIDIA GeForce RTX 5090 GPU (32GB VRAM, CUDA 13.1) with 24 CPU cores and 512GB RAM. SpatialEx was run on the Austrian Scientific Computing (ASC) HPC cluster, using an NVIDIA A40 GPU (48GB VRAM) in a Zen2 node with 16 CPU cores and 256GB RAM.

## Data availability

Newly generated Xenium data will be made available upon publication. The public Xenium v1 public dataset was acquired from Zenodo (DOI: 10.5281/zenodo.18273102).

## Code availability

Newly generated Xenium data will be made available upon publication.

## Acknowledgements

This work was funded by research grants from the Austrian Science Fund (grant number P35937 and KLP2077525) and the LEO Foundation (grant number LF-OC-23-001394) to JG. IO was supported by a DOC fellowship from the Austrian Academy of Science (Grant number 27228). The Vienna Scientific Cluster (Project No. 71839) is gratefully acknowledged for providing computational resources. Funded additionally by the European Union (ERC, AUTODIAL, 101221395). Views and opinions expressed are however those of the author(s) only and do not necessarily reflect those of the European Union or the European Research Council. Neither the European Union nor the granting authority can be held responsible for them.

## Author contributions

**Conceptualization:** RS, JG, PT **Data curation:** JS, RS **Formal analysis:** RS, NS, SG, IO, JS **Funding acquisition:** JG, PT, IO **Methodology:** RS, JG **Software:** RS **Visualization:** RS **Resources:** MS, BS **Supervision:** JG **Writing – original draft:** RS, JG **Writing – review & editing:** all authors

## Ethics declarations

The study protocol was approved by the ethical review board of the Medical University of Vienna (vote 1440/2023).

## Declaration of generative AI use

Generative AI tools (Mistral and chatGPT) were used to assist with code debugging, and to improve language and readability. The authors reviewed and edited the output and take full responsibility for the final content of this publication.

---

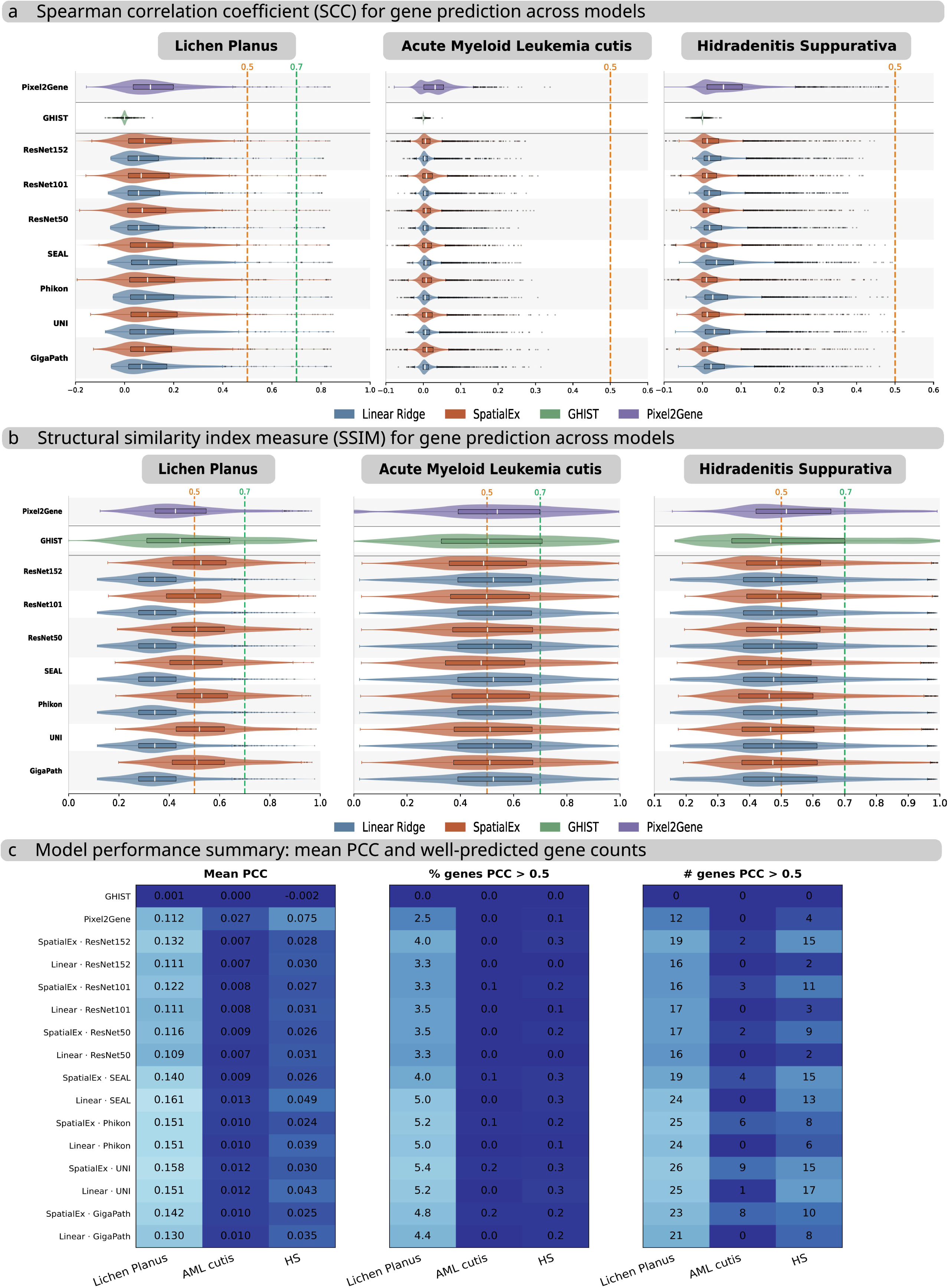

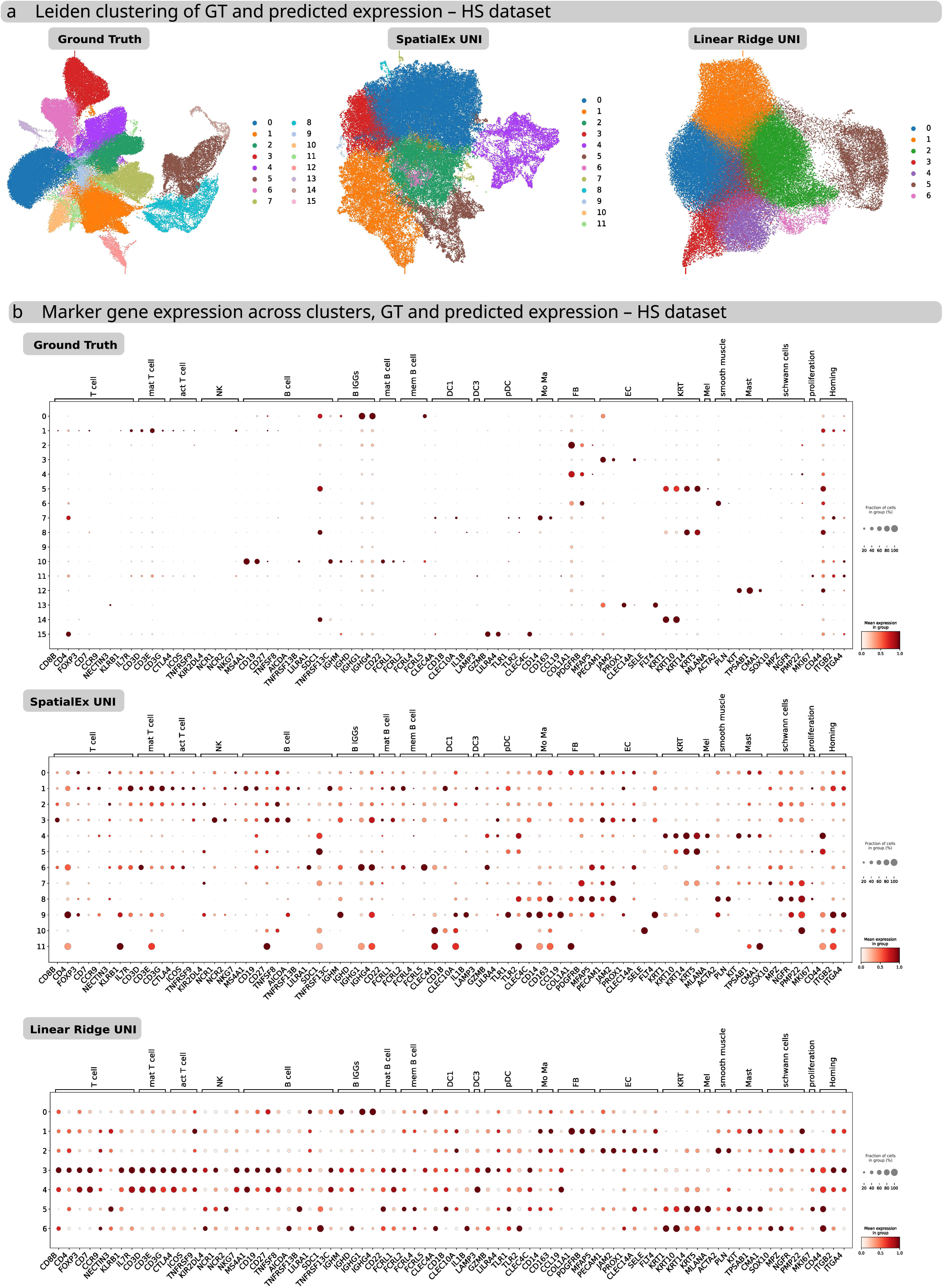

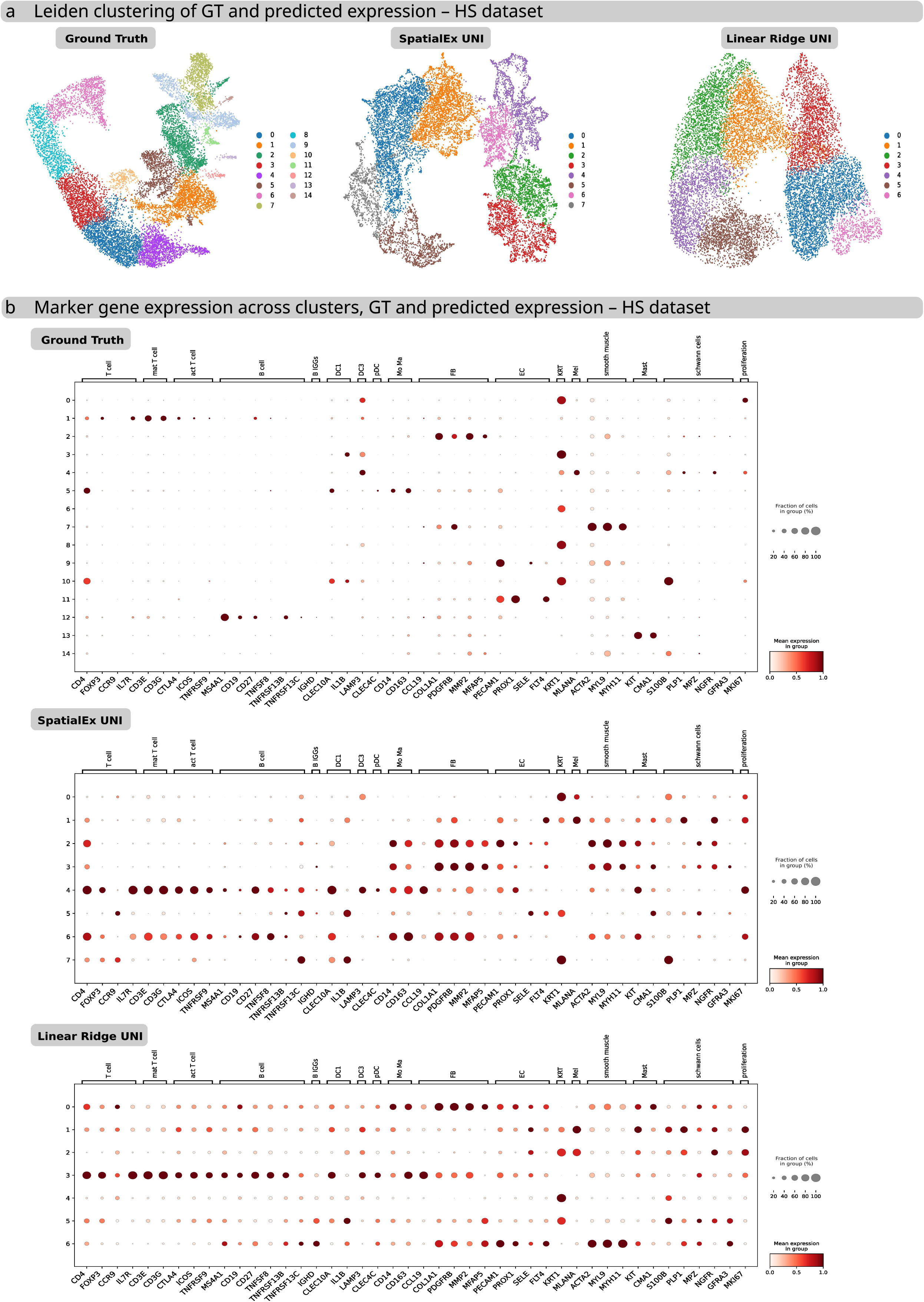

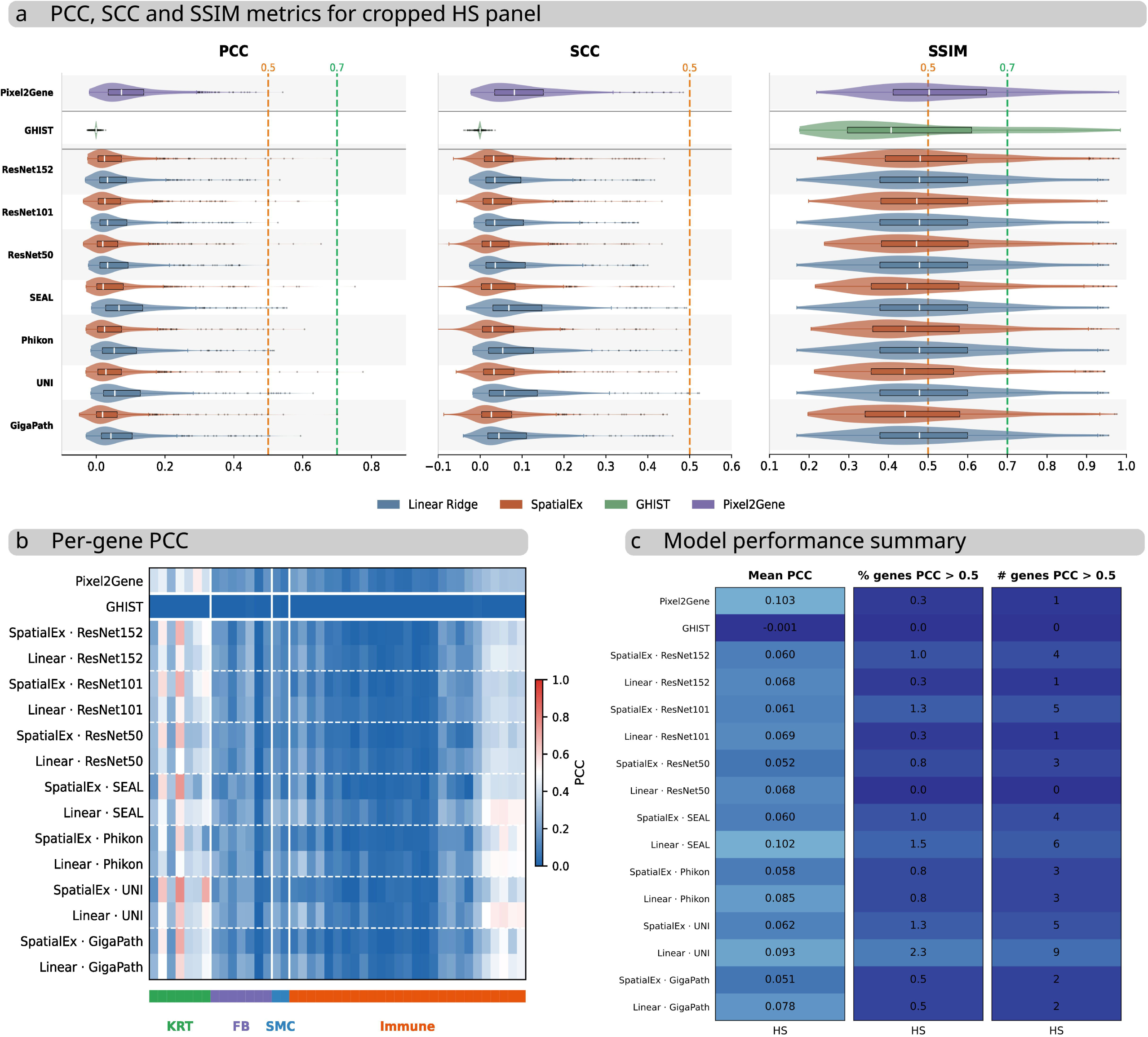

