## Supplementary notes for "Independent benchmark of H&E-based gene expression prediction in skin"

Independent benchmark of H&E-based gene expression prediction in skin - Supplementary Material

### Supplementary Figures

#### Supplementary Figure 1. Additional evaluation metrics and model performance summary.

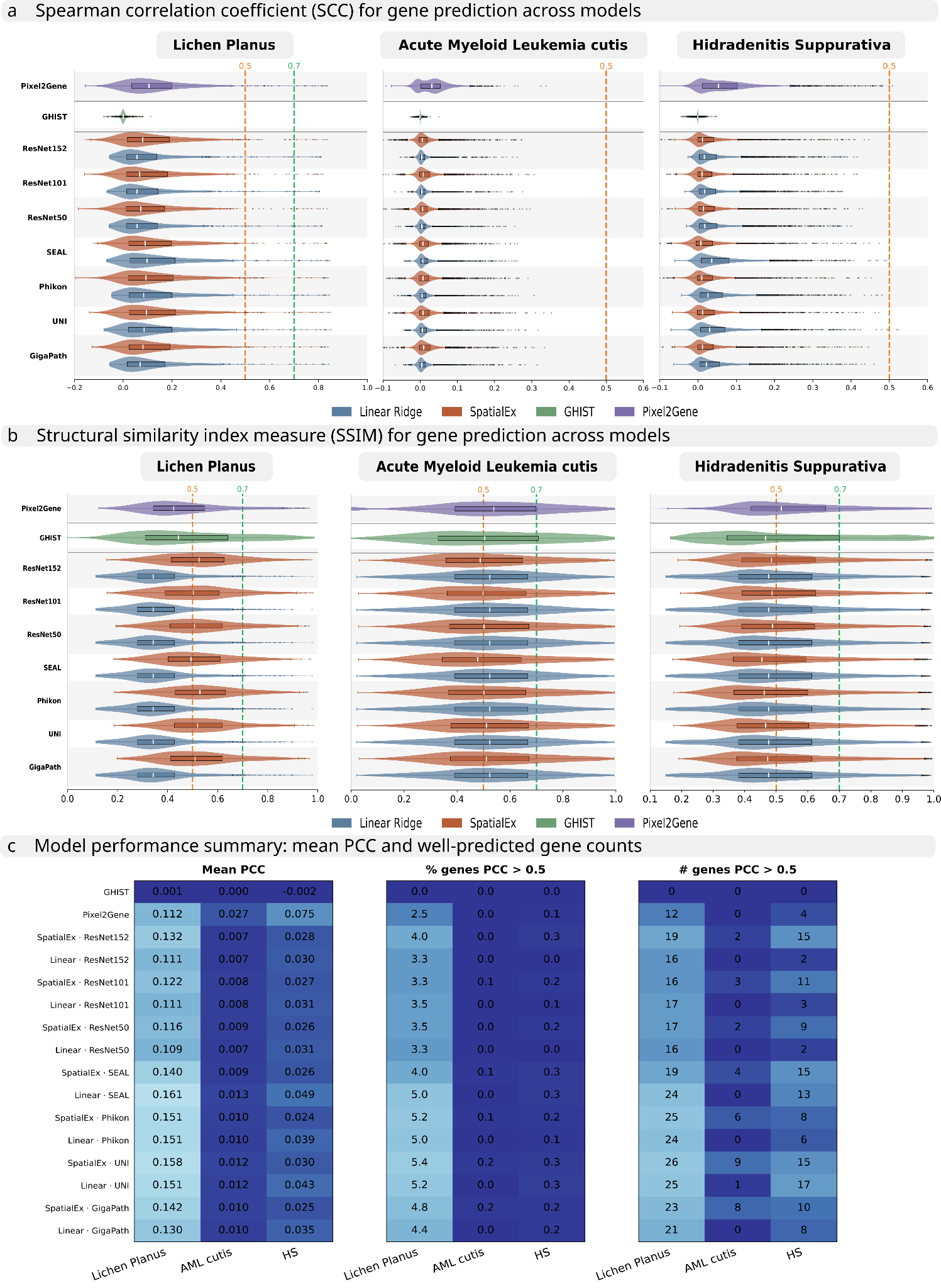

**(a)** Per-gene Spearman correlation coefficient (SCC) per model and dataset. Linear Ridge and SpatialEx per encoder; GHIST, Pixel2Gene separate. Dashed lines mark SCC 0.5, 0.7. **(b)** Per-gene Structural Similarity Index Measure (SSIM) per model and dataset. Linear Ridge and SpatialEx per encoder; GHIST, Pixel2Gene separate. Dashed lines mark SSIM 0.5, 0.7. **(c)** Model performance summary: mean PCC, percentage and number of genes with PCC > 0.5 for all models across all datasets.

**Alt text:** Three-panel performance summary. Panel a: grid of per-gene SCC distributions by model and dataset with dashed lines at 0.5 and 0.7. Panel b: matching grid of per-gene SSIM distributions. Panel c: table/heatmap summarizing mean PCC and the percentage and number of genes with PCC above 0.5 for every model across all datasets.

#### Supplementary Figure 2. Cell-type clustering and annotation for ground truth, SpatialEx UNI, and Linear Ridge UNI in the HS dataset.

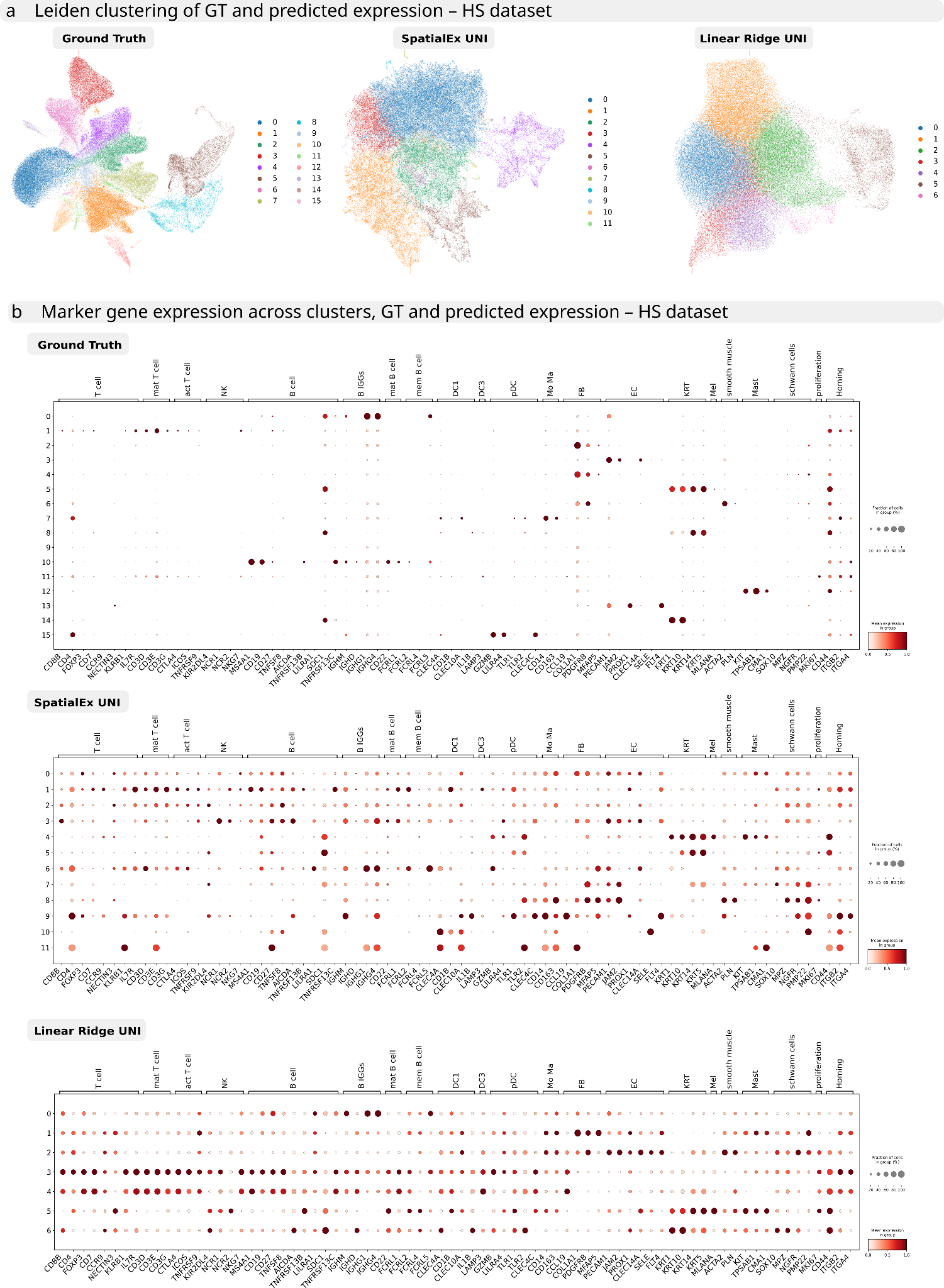

**(a)** UMAP representations of the three datasets: ground truth, SpatialEx UNI prediction, and Linear Ridge UNI prediction; colored by Leiden cluster. **(b)** Marker gene expression across clusters for each of the three datasets, used for cluster annotation.

**Alt text:** Cell-type clustering and annotation in the HS dataset for ground truth, SpatialEx UNI and Linear Ridge UNI. Panel a: three UMAP embeddings colored by Leiden cluster. Panel b: dot/heatmap of marker gene expression across clusters used for annotation.

#### Supplementary Figure 3. Cell-type clustering and annotation for ground truth, SpatialEx UNI, and Linear Ridge UNI in the LP dataset.

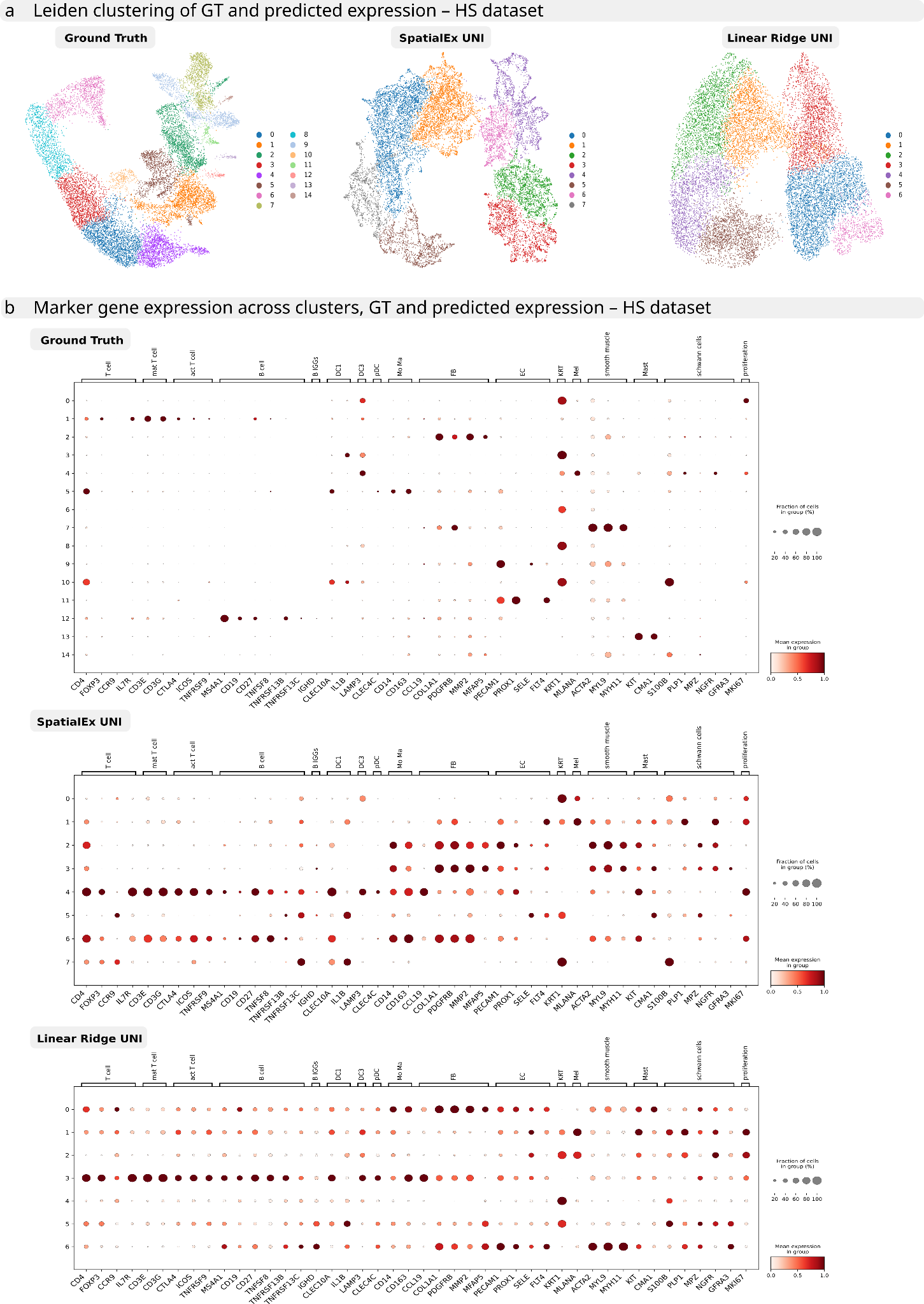

**(a)** UMAP representations of the three datasets: ground truth, SpatialEx UNI prediction, and Linear Ridge UNI prediction; colored by Leiden cluster. **(b)** Marker gene expression across clusters for each of the three datasets, used for cluster annotation.

**Alt text:** Same layout as Supplementary Figure 2 but for the LP dataset: UMAPs colored by Leiden cluster (panel a) and a marker-gene expression heatmap across clusters (panel b).

#### Supplementary Figure 4. Performance of benchmarked models on the HS dataset with a reduced Xenium panel.

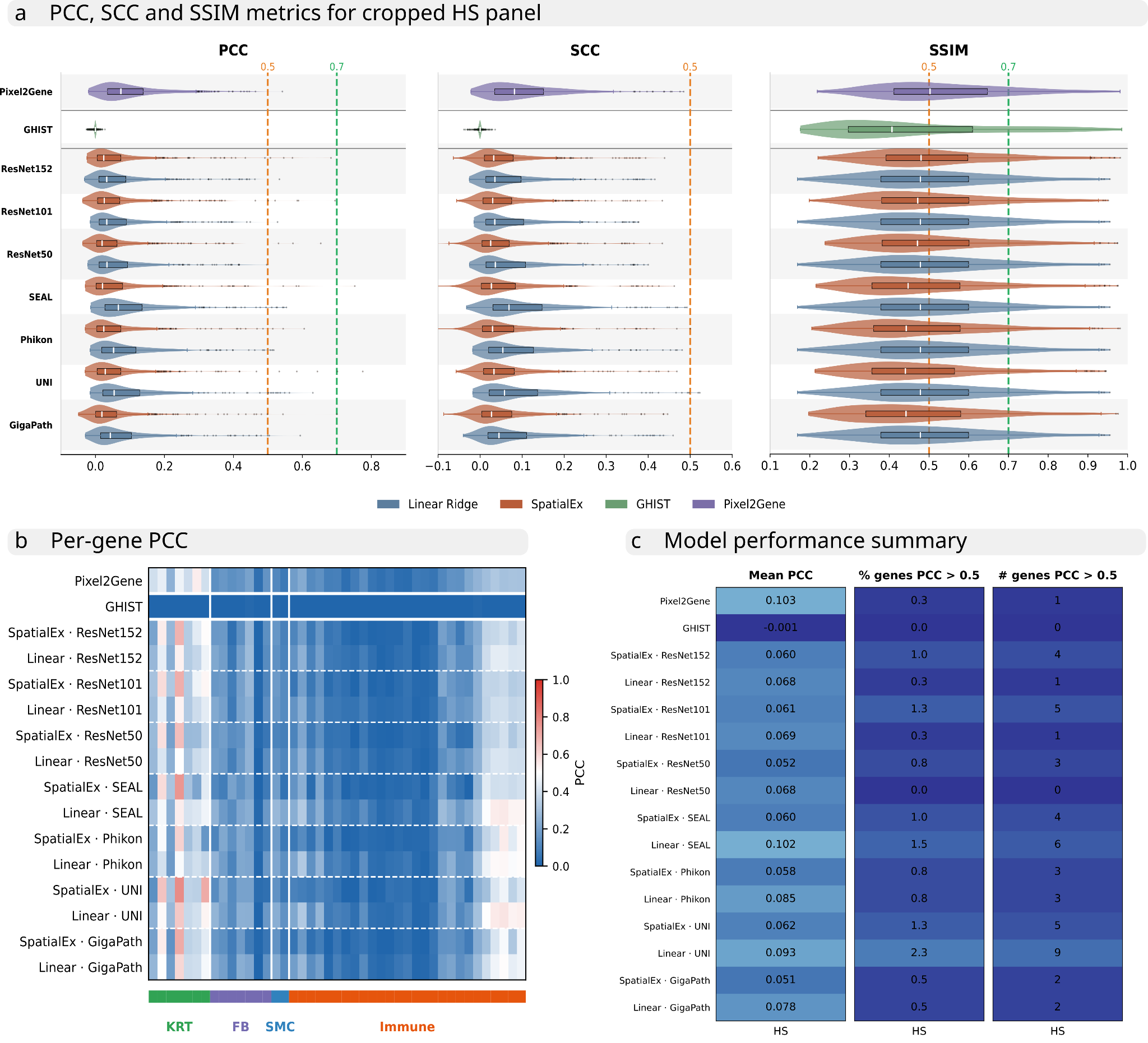

**(a)** PCC, SCC, and SSIM per model. Dashed lines mark 0.5, 0.7. **(b)** Per-gene PCC for cell-type markers grouped (colored bar): keratinocyte (KRT), fibroblast (FB), smooth muscle cell (SMC), immune (Immune). Scale: red 1, blue 0. **(c)** Model performance summary: mean PCC, percentage and number of genes with PCC > 0.5.

**Alt text:** Three-panel figure repeating the benchmark on the HS dataset with a reduced Xenium panel. Panel a: per-gene PCC, SCC and SSIM distributions per model with dashed lines at 0.5 and 0.7. Panel b: heatmap of per-gene PCC for grouped cell-type markers. Panel c: summary table of mean PCC and the percentage and number of genes with PCC above 0.5.

### Supplementary Tables

**Supplementary Table 1. Per-gene Pearson correlation coefficients (PCC) for all models, HS dataset.** PCC between predicted and measured expression for selected cell-type marker genes (rows) across all evaluated models (columns), grouped by cell-type category (KRT, keratinocyte; FB, fibroblast; SMC, smooth muscle cell; Immune). Genes were selected as those reaching PCC > 0.5 in at least one model and dataset, together with additional genes considered relevant for cell-type annotation.

**Alt text:** Data table of per-gene Pearson correlation coefficients between predicted and measured expression in the HS dataset. Rows are selected cell-type marker genes grouped by category (keratinocyte, fibroblast, smooth muscle, immune); columns are the 16 evaluated models.

**Supplementary Table 2. Per-gene Pearson correlation coefficients (PCC) for all models, AML dataset.** PCC between predicted and measured expression for selected cell-type marker genes (rows) across all evaluated models (columns), grouped by cell-type category (KRT, keratinocyte; FB, fibroblast; SMC, smooth muscle cell; Immune). Genes were selected as those reaching PCC > 0.5 in at least one model and dataset, together with additional genes considered relevant for cell-type annotation.

**Alt text:** Same structure as Supplementary Table 1 (per-gene PCC, marker genes by cell-type category as rows, 16 models as columns) for the AML cutis dataset.

**Supplementary Table 3. Per-gene Pearson correlation coefficients (PCC) for all models, LP dataset.** PCC between predicted and measured expression for selected cell-type marker genes (rows) across all evaluated models (columns), grouped by cell-type category (KRT, keratinocyte; FB, fibroblast; SMC, smooth muscle cell; Immune). Genes were selected as those reaching PCC > 0.5 in at least one model and dataset, together with additional genes considered relevant for cell-type annotation.

**Alt text:** Same structure as Supplementary Table 1 (per-gene PCC, marker genes by cell-type category as rows, 16 models as columns) for the LP cutis dataset.

**Supplementary Table 4. Per-gene Pearson correlation coefficients (PCC) for all models, HS dataset, reduced Xenium Panel.** PCC between predicted and measured expression for selected cell-type marker genes (rows) across all evaluated models (columns), grouped by cell-type category (KRT, keratinocyte; FB, fibroblast; SMC, smooth muscle cell; Immune). Genes were selected as those reaching PCC > 0.5 in at least one model and dataset, together with additional genes considered relevant for cell-type annotation.

**Alt text:** Same structure as Supplementary Table 1 (per-gene PCC, marker genes by cell-type category as rows, 16 models as columns) for the HS dataset using the reduced Xenium panel.

### Supplementary Note 1

**Reducing number of genes to the v1 panel does not improve prediction**

To determine whether this difference could be explained by the larger number of target genes in the Xenium 5K panel, we retrained the models on a reduced gene set approximating a Xenium v1 panel. Specifically, we filtered the original set of 5,051 genes to retain only genes present in at least one of the pre-designed Xenium Human Skin Gene Expression Panel or Xenium Human Immuno-Oncology Profiling Panel, resulting in a final set of 397 genes. We then repeated the same analyses as in Fig. 1. The overall patterns of PCC, SCC, and SSIM were highly similar to those observed for the full 5K panel, and we observed no substantial change in model performance (Supplementary Fig. 4a). The same categories of genes, primarily keratinocyte-associated genes and a subset of immunoglobulin genes, remained among the best-predicted genes (Supplementary Fig. 4b). Although mean PCC values were higher in the reduced panel analysis (Supplementary Fig. 4c), this increase was largely driven by the exclusion of lowly expressed and poorly predicted genes. At the level of individual genes, prediction accuracy showed no consistent trend: some genes exhibited modest improvements, whereas others showed reduced performance. For example, CD19 prediction decreased from PCC =~ 0.15 to PCC =~ 0.07 for SpatialEx ResNet152 (Supplementary Table 4). Overall, reducing the number of target genes did not lead to a systematic improvement in prediction accuracy.

### Supplementary Note 2

The SEAL foundation model was the only encoder in our benchmark that was trained on both H&E images and spatial transcriptomics data. However, while the mean PCC for linear ridge SEAL was the highest among all SpatialEx and linear ridge models, the number and proportion of genes achieving PCC > 0.5 were lower than or comparable to those obtained with SpatialEx or linear ridge models based on UNI or Phikon (Supplementary Fig. 1c). This suggests that incorporating spatial transcriptomics data into the foundation model's pretraining did not improve prediction performance for skin tissue
